# Extracellular Vesicle Gene Expression Enables Sensitive Detection of Colorectal Neoplasia

**DOI:** 10.1101/2025.08.14.670394

**Authors:** Sivakumar Gowrisankar, J. Christian J. Ray, Sinead Nguyen, Christopher J. Benway, Shuran Xing, Allan George, Dulaney L. Miller, Kyle Manning, Yang Yang, Jeff Cole, Emily Mitsock, Devin Perez, Luanna Dealmeida, Julia Doo, Pawan Kumar Upadhyay, Yiyuan Yao, Joseph M. Johnson, Brian C. Haynes, Sudipto K. Chakrabortty, Johan K. Skog

## Abstract

**BACKGROUND & AIMS:** Extracellular vesicles (EVs), including exosomes, are emerging as promising carriers of disease-specific biomarkers due to their molecular cargo reflective of cellular origin. While cell-free DNA (cfDNA) methylation assays have recently been developed for colorectal cancer (CRC) screening and perform well for cancer detection, they show limited sensitivity for advanced adenomas (AA), a key precursor in the CRC pathway. To address this gap, we sought to determine whether other blood-based analytes, specifically EV-derived long and small RNA transcriptomes and EV proteomes, could improve detection of AA and early-stage CRC. In a prospective cohort, we performed a head-to-head comparison of EV transcriptomics, EV proteomics, and their combinations against cfDNA methylation, all measured from the same patient cohort, enabling a direct performance benchmark and identification of the most promising modality for larger-scale CRC screening studies.

**METHODS:** We prospectively collected pre-colonoscopy plasma samples from 220 participants across three clinical sites. EVs were isolated and profiled using long RNA-seq, small RNA-seq, and Olink-based proteomics. cfDNA was analyzed for methylation patterns. Analyses were conducted according to a statistical analysis plan pre-specified before unblinding. Machine learning models were developed under nested cross-validation to evaluate sensitivity for detecting AA and CRC at a fixed specificity of 91%, consistent with clinical screening benchmarks.

**RESULTS:** EV-derived gene expression on long RNA demonstrated the highest sensitivity for detecting AA as well as CRC: 54.8% (95% CI, 26.4%–75.6%) for AA and 94.1% (95% CI, 79.2%–100%) for CRC. This outperformed cfDNA methylation (33.6% [95% CI, 9.7%–60.3%] for AA, 81.3% [95% CI, 64.6%– 93.9%] for CRC) and other EV-based modalities. In addition, for small RNA the sensitivities were 42.6% (95% CI, 35.9%-47.4%) for AA, and 78.9% (95% CI, 69.2%-83.0%) for CRC, while for proteomics the sensitivities were 30.0% (95% CI, 13.9%-40.0%) for AA, and 64.2% (95% CI, 43.6%-87.2%) for CRC. Transcriptomic profiles revealed progressive enrichment of hallmarks of cancer pathways, including apoptosis and epithelial-mesenchymal transition, across disease stages. Multiomic integration did not improve performance beyond EV transcriptomics alone.

**CONCLUSIONS:** By directly comparing multiple EV-based and cfDNA analytes within the same patient cohort, we found that EV transcriptomics delivers the strongest diagnostic performance for both AA and CRC. This rigorous benchmarking approach allows clear prioritization of the most promising modality guiding the design of larger validation studies and accelerating development of next-generation, blood-based CRC screening tools.

## Introduction

Colorectal cancer (CRC) represents a major global health challenge, consistently ranking as the third most diagnosed malignancy, and the second leading cause of cancer-related mortality worldwide.^1^ Screening and early detection remain one of the most effective strategies for reducing mortality, particularly for malignancies arising in tissues that are not easily accessible for routine surveillance. CRC often develops from precancerous polyps that can be detectable during a colonoscopy, making it one of the few cancers that is truly preventable through early intervention^2^, Despite improvements in noninvasive options like Cologuard, overall colorectal cancer (CRC) screening participation remains around 60% in the U.S, well below the National Colorectal Cancer Roundtable’s 80% goal. on-invasive approaches, This underscores a growing need for more acceptable blood-based screening alternatives.^3^ In CRC, blood-based assays incorporating cell-free DNA (cfDNA), proteins, and clinical covariates have demonstrated moderate success in detecting Stage I or later disease.^4^ However, sensitivity for identifying pre-cancerous lesions such as advanced adenomas remains very limited.^5^

Tissue-based studies, including single-cell analyses, have revealed extensive molecular alterations in advanced adenomas and CRC, including oncogenic mutations and dysregulated gene expression.^6^ Despite this, the low abundance of circulating cfDNA in plasma may constrain the sensitivity of blood-based tests for early-stage disease, in contrast to stool-based assays which are more proximal to the tumor site.

Extracellular vesicles (EVs), including exosomes, are released by nearly all cell types and carry molecular cargo – including RNA, DNA, and proteins – that reflects the physiological or pathological state of their cell of origin. Their stability and ability to encapsulate tumor-specific biomarkers make them a promising platform for noninvasive early detection of CRC.^7^ We hypothesized that cfDNA would serve as the baseline for performance in detecting biological processes underlying CRC development, including pre-cancerous stages, and that RNA and protein cargo from circulating extracellular vesicles (EVs) could complement limitations of cfDNA-based liquid biopsies. To test this, we conducted a multiomics feasibility study using plasma samples collected from a screening population prior to colonoscopy (Figure 1a). Plasma was processed to isolate cfDNA for methylation analysis and EVs for long RNA-seq, small RNA-seq, and Olink-based proteomics.

**Figure 1.**
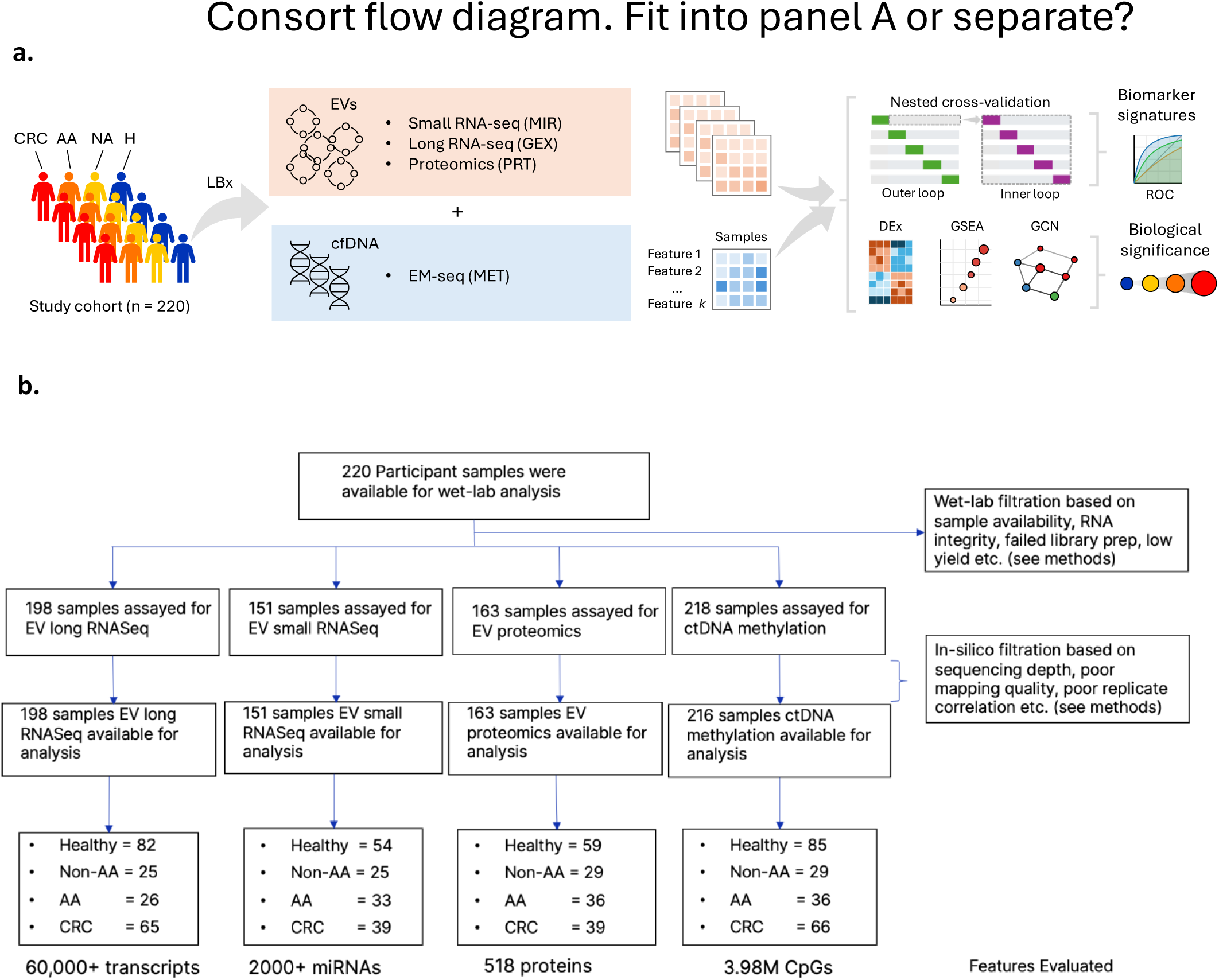
Design of a multiomic study to assess the capabilities of EV-derived signals for detection of colorectal cancer progression. **a.** Plasma samples were collected prior to colonoscopy and associated pathology reports were collected after colonoscopy. Samples were divided for isolation of EVs and cfDNA and subjected to multiomic analysis. **b.** Consort flow diagram for the multiomic study.

Following data generation, multiomic profiles were transformed into feature-by-sample matrices and analyzed using machine learning models incorporating robust feature selection and sensitivity estimation. These models were designed to predict the presence of advanced adenomas (AAs) and colorectal cancer. Our findings demonstrate that EV-derived gene expression alone provides enhanced sensitivity, identifies biologically relevant features, and mirrors known molecular pathways involved in CRC progression. These results underscore the feasibility of a larger validation study to establish EV-derived gene expression, and other EV-biomarkers for the early and accurate detection of colorectal neoplasia.

## Methods

Materials and Methods are provided as part of supplementary materials.

## Results

### Study Population

A total of 220 participants were included: 87 healthy controls, 29 with non-advanced adenomas (Non-AA), 36 with advanced adenomas (AA), and 68 with colorectal cancer (CRC). Median age increased with disease severity (Healthy: 51 years; CRC: 64 years). The cohort was predominantly Caucasian (92.2%), with balanced gender representation (51.6% female). CRC staging data were available for 59 of the 68 cancer cases. Stage III was the most prevalent (48.5%), followed by Stage IV (20.6%), Stage I (8.8%), and Stage II (8.8%). All CRC samples irrespective of the stage annotations were included in the analysis, as the primary model performance was evaluated for the screening ability of CRC, irrespective of stage. Importantly, the study included early-stage cases, enabling evaluation of biomarker performance in Stage I and II disease, which is critical for screening applications. Limited serratedness and tubulovillous status data were available for AA and Non-AA patients (Table 1). Of the 220 total participants enrolled, 198 participants for EV Gene Expression, 151 participants EV small RNA, 163 participants for EV proteomics, and 216 participants for ctDNA methylation could be fully evaluated. Sample loss at the wet-lab level was due to a combination of factors including sample availability (volume), RNA integrity, library prep QC, and low library yield. Filtration of samples at the *in silico* step was based on sequencing depth and poor spike-in replicate correlation among other factors (Figure 1b; see Methods).

**Table 1.** Demographics.

able
|  | Healthy (N=87) | Non-Advanced Adenoma (N=29) | Advanced Adenoma (N=36) | Colorectal Cancer (N=68) | Total (N=220) |
| --- | --- | --- | --- | --- | --- |
| Age, median (IQR) | 52.0 (42.0-61.0) | 51.0 (47.0-64.0) | 58.5 (48.8-65.2) | 64.0 (55.0-70.0) | 57.0 (48.0-66.0) |
| Height (cm) , median (IQR) | 172.0 (165.0-179.0) | 168.0 (162.0-175.0) | 172.5 (164.8-176.5) | 168.0 (165.0-176.0) | 170.5 (165.0-178.0) |
| Weight (kg) , median (IQR) | 80.0 (70.5-94.0) | 84.0 (75.0-90.0) | 79.5 (72.8-87.5) | 74.0 (67.2-80.0) | 79.0 (70.8-90.0) |
| Race, n (%) |  |  |  |  |  |
| Asian | 2 (2.3%) | 0 (0.0%) | 0 (0.0%) | 0 (0.0%) | 2 (0.9%) |
| Black/African American | 7 (8.0%) | 0 (0.0%) | 0 (0.0%) | 0 (0.0%) | 7 (3.2%) |
| Caucasian | 75 (86.2%) | 29 (100.0%) | 36 (100.0%) | 68 (100.0%) | 208 (94.5%) |
| Multiracial | 1 (1.1%) | 0 (0.0%) | 0 (0.0%) | 0 (0.0%) | 1 (0.5%) |
| Gender, n (%) |  |  |  |  |  |
| Female | 46 (52.9%) | 20 (69.0%) | 15 (41.7%) | 33 (48.5%) | 114.(51.8%) |
| Male | 39 (44.8%) | 9 (31.0%) | 21 (58.3%) | 35 (51.5%) | 104 (47.3%) |
| Smoking Status, n (%) |  |  |  |  |  |
| Current | 4 (4.6%) | 5 (17.2%) | 6.(16.7%) | 0 (0.0%) | 15 (6.8%) |
| Former | 8 (9.2%) | 7 (24.1%) | 10 (27.8%) | 6 (8.8%) | 31 (14.1%) |
| Never | 27 (31.0%) | 17 (58.6%) | 20 (55.6%) | 14 (20.6%) | 78 (35.5%) |
| Alcohol Status, n (%) |  |  |  |  |  |
| Current | 17 (19.5%) | 10 (34.5%) | 11 (30.6%) | 5 (7.4%) | 43 (19.5%) |
| Former | 1 (1.1%) | 0 (0.0%) | 0 (0.0%) | 0 (0.0%) | 1 (0.5%) |
| Never | 21 (24.1%) | 19 (65.5%) | 25 (69.4%) | 15 (22.1%) | 80 (36.4%) |
| Family History, n (%) |  |  |  |  |  |
| False | 34 (39.1%) | 26 (89.7%) | 31 (86.1%) | 17 (25.0%) | 108 (49.1%) |
| True | 5 (5.7%) | 3 (10.3%) | 5 (13.9%) | 3 (4.4%) | 16 (7.3%) |
| Tubulovillous, n (%) |  |  |  |  |  |
| False | 87 (100.0%) | 28 (96.6%) | 30 (83.3%) | 68 (100.0%) | 213 (96.8%) |
| True | 0 (0.0%) | 1 (3.4%) | 6 (16.7%) | 0 (0.0%) | 7 (3.2%) |
| Serratedness, n (%) |  |  |  |  |  |
| False | 87 (100.0%) | 25 (86.2%) | 31 (86.1%) | 68 (100.0%) | 211 (95.9%) |
| True | 0 (0.0%) | 4 (13.8%) | 5 (13.9%) | 0 (0.0%) | 9 (4.1%) |
| Advanced Precancerous Lesions, n (%) |  |  |  |  |  |
| False | 87 (100.0%) | 29 (100.0%) | 21 (58.3%) | 68 (100.0%) | 205 (93.2%) |
| True | 0 (0.0%) | 0 (0.0%) | 15 (41.7%) | 0 (0.0%) | 15 (6.8%) |
| Stage, n (%) |  |  |  |  |  |
| 1 | 0 (0.0%) | 0 (0.0%) | 0 (0.0%) | 6 (8.8%) | 6 (2.7%) |
| 2 | 0 (0.0%) | 0 (0.0%) | 0 (0.0%) | 6 (8.8%) | 6 (2.7%) |
| 3 | 0 (0.0%) | 0 (0.0%) | 0 (0.0%) | 33 (48.5%) | 33 (15.0%) |
| 4 | 0 (0.0%) | 0 (0.0%) | 0 (0.0%) | 14 (20.6%) | 14 (6.4%) |

### Omics Profiles Indicate Better Signal with EV-derived Gene Expression

To characterize molecular signatures associated with colorectal neoplasia, we performed comprehensive multiomic profiling on plasma-derived EVs and cfDNA. The modalities included RNA-seq of long transcripts for gene expression, small RNA-seq for miRNAs, Olink-based proteomics, and methylation profiling of cfDNA. First, an unsupervised analysis was conducted to explore data structure and predictive potential.

To investigate the underlying structure of the multiomic data, we performed unsupervised principal component analysis (PCA) on each data type for healthy, non-advanced adenoma (non-AA), advanced adenoma (AA), and colorectal cancer (CRC) samples. The analysis of methylation features revealed a good separation between healthy and CRC samples along the first principal component (Figure 2a). While a few CRC samples clustered with the adenoma and healthy groups, a subset of CRC samples formed a distinct, separate cluster. A similar separation was observed in the PCA of gene expression features (Figure 2d). In contrast, the PCA of EV proteins (Figure 2b) and EV miRNAs (Figure 2c) did not show clear clustering by disease state, with samples from all conditions exhibiting considerable overlap.

**Figure 2.**
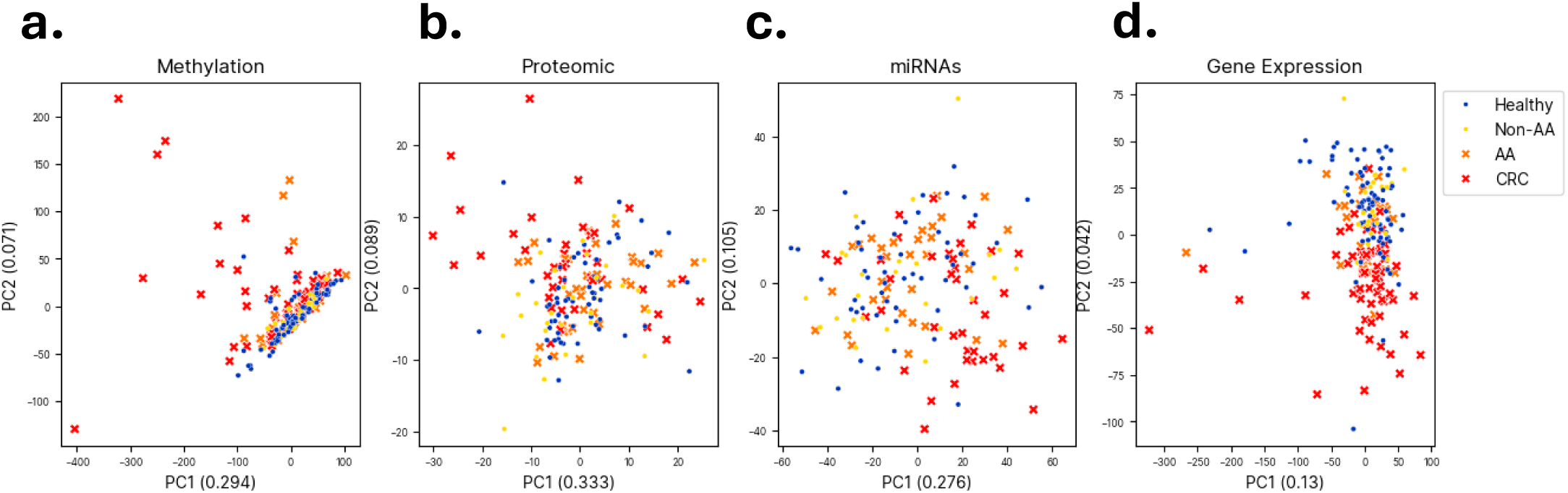
Unsupervised principal component analysis (PCA) of plasma multiomics in stages of colorectal cancer development. Samples are colored by condition as indicated in the legend. The fractional variance contribution of each principal component is in parentheses. **a.** PCA of methylation features. **b.** PCA of EV proteins. **c.** PCA of EV miRNAs. **d.** PCA of gene expression features.

Next, beyond the separation observed in PCA plots, we quantified the magnitude and fold change for each feature across omics layers, comparing Healthy + Non-AA and AA + CRC groups. Probability density estimates revealed broader distributions for EV gene expression and miRNAs, indicating a greater dynamic range and biological variability. In contrast, cfDNA methylation and EV proteomics exhibited narrower distributions, suggesting more constrained dynamic ranges (Figure 3a). Quantile-quantile (Q-Q) plots further supported these findings, showing marked deviations from normality— particularly in gene expression and methylation datasets—highlighting the presence of outlier features with strong discriminatory potential (Figure 3c). It must be noted that the methylation data set has a strong tail towards the left but also a heavy skew towards the right, indicating more hypermethylated than hypomethylated signal. Both miRNA and gene expression show heavy-tailed distributions on both the right and left, indicating the presence of outliers for both up and down-regulated gene ends than expected. To comprehensively assess discriminatory power, we calculated the mean and 95% confidence intervals of absolute Log2 fold change (Log2FC) for each feature across all samples within each omics layer. This analysis complements the density plots by highlighting the central moment of the distributions, revealing a progressive increase in average effect size from cfDNA methylation to EV gene expression. Notably, EV-derived miRNAs demonstrated a relatively small confidence interval and large effect size, whereas EV gene expression showed the second-largest average effect size accompanied by a small confidence interval (Figure 3b). The strong signal within the EV RNA-seq data was further confirmed by differential expression analysis, which revealed a substantial number of significantly altered genes that increased with disease severity (Table 2). For instance, when comparing CRC patients to the control group (Healthy + Non-AA), we identified 5,709 differentially expressed genes at a false discovery rate (FDR) of 0.05. This number remained high even at a stringent FDR of 0.01, with 4,042 significant genes. In contrast, the comparison between only AA patients and controls yielded just 17 significant genes at an FDR of 0.05. This large and robust signal in both pre-cancerous and cancer stages of disease suggests that both EV gene expression and miRNA signatures contain high-quality information that could be leveraged for building powerful diagnostic and prognostic models.

**Figure 3.**
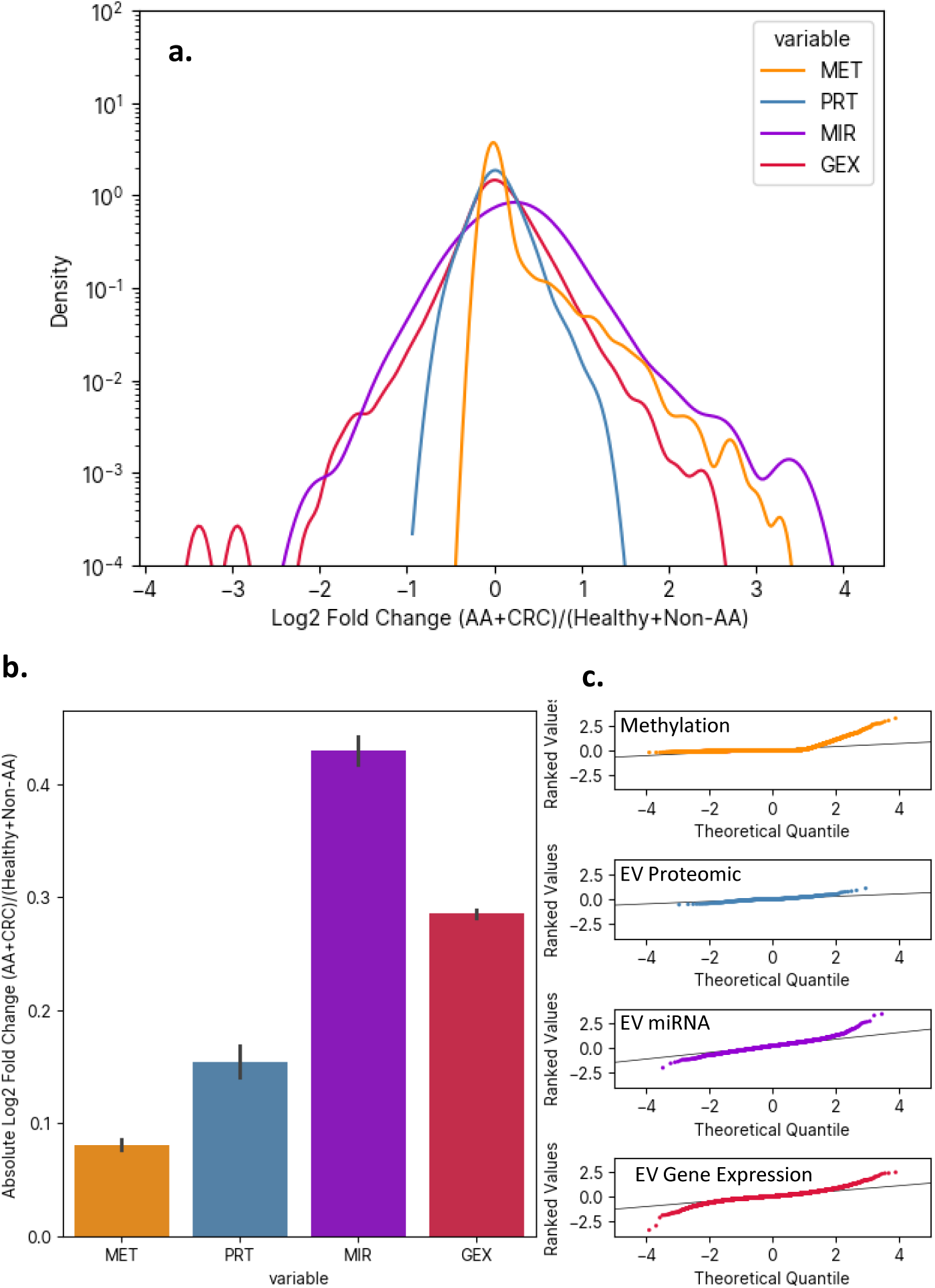
Distribution of effect sizes across omics. **a.** Density of log2 fold change between the positive and negative groups. **b.** Central moment +/− 95% CI of fold changes between the positive group (AA+CRC) and the negative group (Healthy + Non-AA). MET, methylation; PRT, proteomic; MIR, miRNAs; GEX, long RNA gene expression (mRNAs and long-noncoding RNA). **c.** Quantile-quantile plots for each omic reveals divergence from the fitted normal distribution (black line).

**Table 2.**
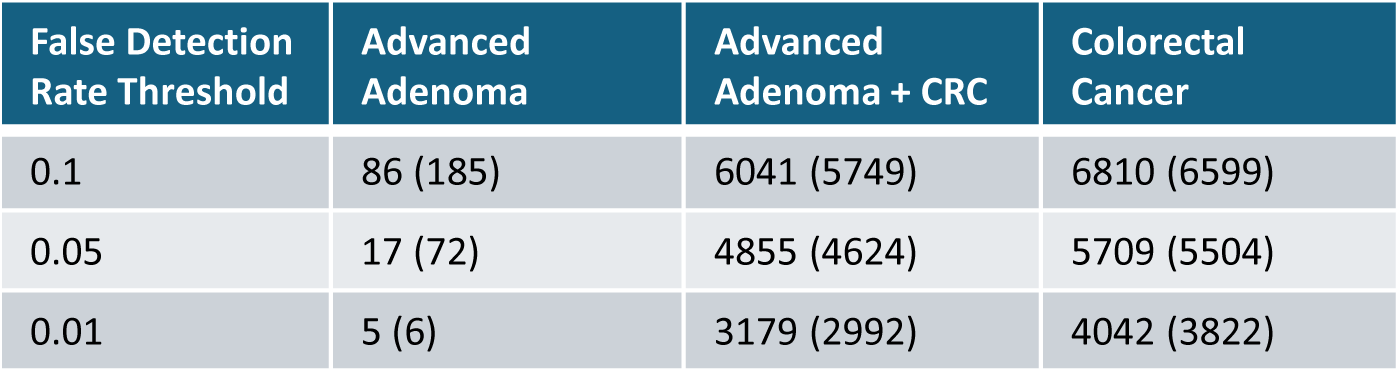
The number of significantly differentially expressed genes from EV RNA-seq. Parentheses: comparisons that exclude non-advanced adenomas.

### Biological Relevance of EV Gene Expression in CRC Progression

To investigate the biological relevance of EV-derived gene expression in CRC progression, we performed pathway-level analyses using gene set variation analysis (GSVA) and weighted gene co-expression network analysis (WGCNA).^8,9^ These approaches enabled us to take an unsupervised view into sample groups and assess coordinated transcriptional changes across disease stages.

GSVA enrichment scores were calculated for hallmark cancer pathways, revealing progressive shifts in pathway activity from healthy individuals to those with advanced adenomas and CRC. As shown in Figure 4a, the apoptosis and epithelial-mesenchymal transition (EMT) pathways exhibited a clear increase in enrichment scores across disease stages. This trend is consistent with the known role of apoptosis dysregulation in tumor initiation and progression.^10^ Similarly, EMT is a key driver of tumor invasion and metastasis, and its detection in circulating EVs suggests that these vesicles capture transcriptional programs linked to cellular plasticity and metastatic potential.^11^ In healthy samples, pathway activity was low, while advanced adenoma and CRC samples showed elevated scores, suggesting that EVs carry transcripts reflective of cellular stress and programmed cell death mechanisms. This progressive increase in pathway activity was not limited to Apoptosis and EMT. A comprehensive analysis across all 50 hallmark gene sets revealed that the majority showed a statistically significant, stepwise increase in enrichment from healthy controls to advanced adenoma and finally to CRC (ANOVA FDR < 0.05; Figure S5). This pattern was particularly evident in pathways fundamental to cancer biology, including those governing cell cycle progression (e.g., G2M Checkpoint, E2F Targets), proliferation (e.g., MYC Targets V1 and V2), and responses to genomic instability (e.g., DNA Repair).^12^ The widespread activation of these oncogenic signaling pathways, reflected in the circulating EV transcriptome, underscores the potential of this approach to provide a systemic and non-invasive snapshot of the tumor’s biological state. The specificity of this signature is highlighted by the fact that several pathways, such as Bile Acid Metabolism and Coagulation, remained unchanged across the disease continuum, indicating a targeted alteration of cancer-related processes rather than a non-specific global change.

**Figure 4.**
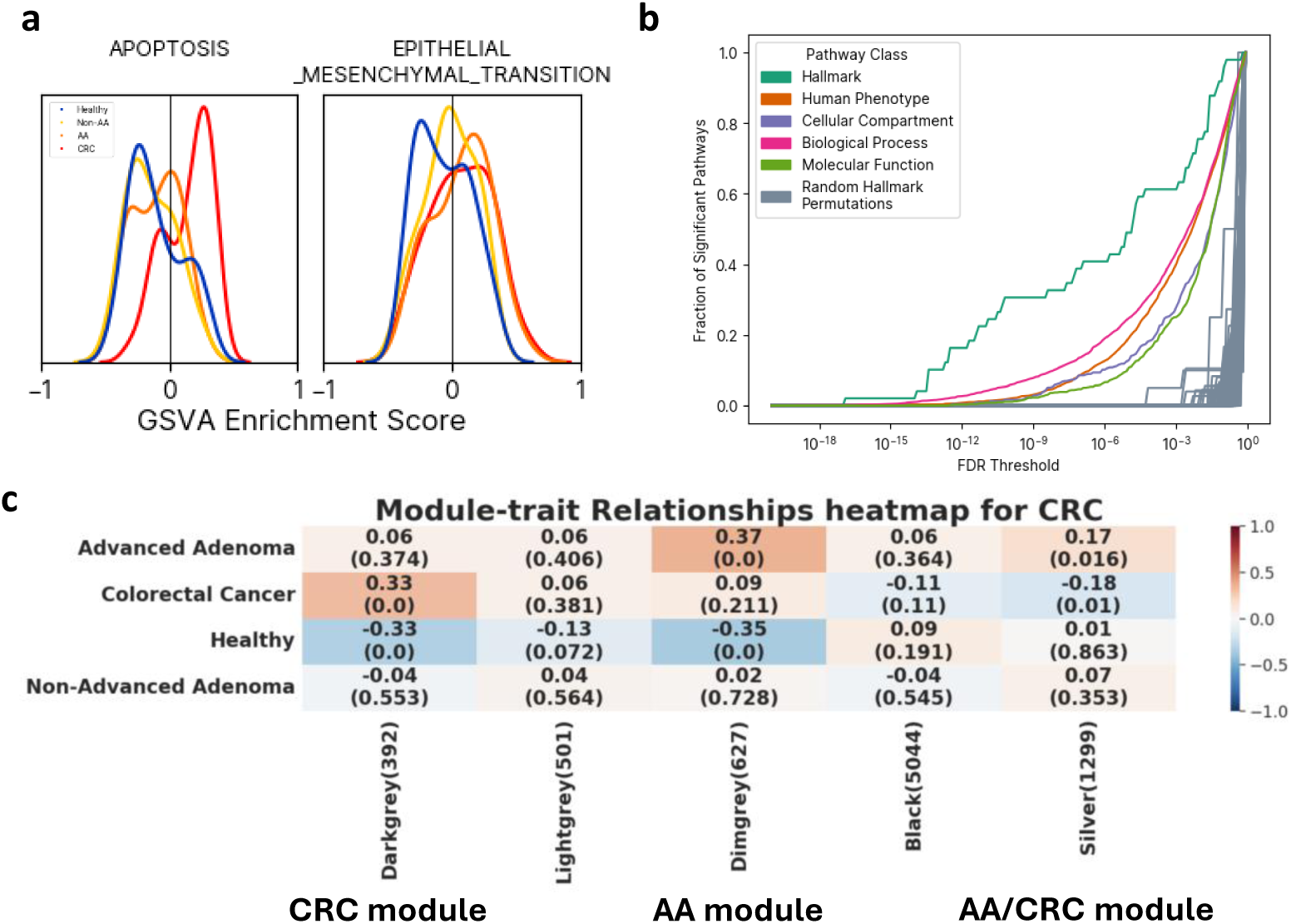
Gene expression in plasma-derived EVs at the pathway level in progressive stages of colorectal cancer development. **a.** Example Hallmark pathway enrichment score distributions for two key cancer pathways from gene set variation analysis. **b.** ANOVA F-test significant pathway numbers for a scan of the false detection rate (FDR) threshold reveals more enriched pathways than a randomly permuted set of Hallmark pathway scores (gray lines; N=100 permutations). **c.** WGCNA predicts modules in EV gene expression of colorectal cancer progression, with each disease/module populated with correlation coefficient of eigen genes(p value in parentheses). Modules with significant correlations are named (i.e., CRC, AA, and AA/CRC modules). Other modules (Lightgrey, Black) are not used for further analysis.

To rigorously evaluate the statistical significance of observed pathway alterations, we implemented a multi-step analytical approach. First, we employed an ANOVA F-test to compare pathway enrichment scores across our four study cohorts: healthy, Non-AA, AA, and CRC. Recognizing that testing thousands of pathways simultaneously inflates the probability of chance discoveries, we corrected for multiple hypotheses by controlling the False Discovery Rate (FDR) using the Benjamini-Hochberg^13^ method. To establish a robust, empirically derived null model, we then performed permutation testing. This involved creating 100 shuffled datasets where the sample phenotype labels were randomly reassigned, thereby breaking the true biological relationships while preserving the underlying data structure. By analyzing these permuted datasets, we generated a realistic baseline for the number of significant pathways that could arise purely by chance. As illustrated in Figure 4b, we observed a distinct and substantial separation between the fraction of significant pathways in our empirical data (colored lines) and that of the permuted null distribution (gray area). This gap persisted across the entire spectrum of FDR thresholds, demonstrating that the number of altered pathways in our dataset far exceeds what would be expected by random chance, even at the most stringent levels of statistical significance. Therefore, we conclude that the identified pathway dysregulations are not statistical artifacts but represent a robust and non-random biological signal that is fundamentally characteristic of colorectal cancer initiation and progression.

We next applied WGCNA analysis to identify modules of co-expressed genes associated with disease states.^9^ In brief, WGCNA identifies groups of genes, called modules, that have highly correlated expression patterns across a set of samples. Figure 4c presents a heatmap that visualizes the relationship between specific WGCNA modules and the different stages of colorectal cancer progression. It highlights three modules – which we labeled CRC, AA, and AA/CRC *post hoc* – with distinct correlations to CRC progression. Each cell shows the correlation coefficient (top number) and its statistical significance (p-value, in parentheses), with red indicating a positive correlation and blue indicating a negative correlation. The CRC and AA modules show very similar patterns in that both have a strong and statistically significant negative correlation with the Healthy state (CRC module correlation coefficient: −0.33, p<1×10^−5^ AA module correlation coefficient: −0.35, p<1×10^−5^). However, while the CRC module has positive correlation with CRC (correlation coefficient: 0.33, p<1×10^−5^), the AA module has positive correlation with AA (correlation coefficient: 0.37, p<1×10^−5^). The AA/CRC module, however, presents a more distinct signature that distinguishes AA and CRC patients. It is not only negatively correlated with CRC state (−0.18, p=0.01) but is also positively correlated with AA (0.17, p=0.016). This opposing relationship suggests that the 1,299 genes in the AA/CRC module are particularly relevant to cancer progression, as their expression levels increase specifically with the onset of malignant disease.

To further explore the biological underpinnings of the EV-derived gene expression signatures associated with colorectal cancer in these modules, we performed gene ontology (GO) enrichment analysis using Python modules^14^ referencing MSigdb.^15^ In the CRC module, GO enrichment analysis revealed pathways central to tumor biology, including cell cycle regulation, metabolic reprogramming, and cell adhesion (Figure S1). These findings are consistent with the proliferative and immunomodulatory phenotype of advanced CRC.^16,17^ The presence of these signatures in EV-derived RNA supports the hypothesis that circulating vesicles reflect active tumor processes and can serve as a non-invasive window into disease biology. In the AA module, the enrichment of pathways related to the innate immune system, points to a state of chronic inflammation within the advanced adenoma microenvironment. This finding is consistent with the well-established role of inflammation as a key driver in the adenoma-to-carcinoma sequence (Figure S2).^18,19^ These features suggest that EVs may carry early indicators of tissue remodeling and immune activation, offering potential for detecting pre-cancerous lesions before malignant transformation.^12,20,21^ In the AA/CRC module, enriched GO terms indicate that malignant progression is associated with the suppression of pathways critical for tissue homeostasis and immune function (Figure S3). Taken together, these results profile the transition from AA to CRC as a process involving the loss of cellular specialization, the breakdown of tissue structure, and the suppression of active immune surveillance.^12^ The presence of these signals in EVs suggests that this module captures intermediate biology relevant to early detection and disease monitoring.

Together, these findings demonstrate that EV-derived gene expression captures coordinated, stage-specific transcriptional programs relevant to CRC development. The integration of GSVA, FDR-based enrichment, and WGCNA provides a systems-level view of disease biology and supports the use of EVs as a non-invasive source of pathway-level biomarkers. These results are visualized in Figure 4, which summarizes the progression-associated changes in EV transcriptomic profiles.

### Deconvolution of Plasma EVs Reveals a Prominent Colon Signature

To investigate the tissue origins of circulating EV-RNA and determine if their proportional contributions change during colorectal cancer (CRC) progression, we performed computational deconvolution. Using the DeconRNASeq algorithm^22^ and reference transcriptomes from the GTEx consortium^23^, we estimated the relative contribution of 12 major tissue types to the plasma EV-RNA pool across our healthy, Non-AA, AA, and CRC cohorts (Figure S4).

Our analysis revealed that hematopoietic tissues were the primary source of plasma EV-RNA across all patient groups. As expected, lymphocytes and whole blood represented the two largest contributors. Notably, the colon was identified as the third most abundant contributor, with a median contribution of approximately 15% across all cohorts. This substantial signal underscores the significant and consistent input of the gastrointestinal system to the systemic circulating EV pool. Other tissues, such as the artery, spleen, and liver, showed modest contributions, while tissues like the esophagus and small intestine represented only a minor fraction of the total EV-RNA signature.

To statistically assess whether the tissue-of-origin profile was altered by disease status, we applied a one-way ANOVA for each of the 12 tissue types, followed by a False Discovery Rate (FDR) correction for multiple comparisons. Despite the high contribution from the colon, its proportional representation did not differ significantly between the disease groups. In fact, the analysis revealed that only the contribution from whole blood was significantly altered across the cohorts (Padj = 0.021). This was characterized by a discernible decrease in the EV-RNA contribution from blood in the colorectal cancer group compared to the healthy cohort. No other tissue type demonstrated a statistically significant change in its contribution after correction.

### Evaluation of Detection Accuracy for Advanced Adenoma and Colorectal Cancer

Supervised machine learning models were trained using nested cross-validation to ensure robust performance estimation. Feature-by-sample matrices were constructed for each omic (i.e., EV gene expression, EV proteomics, EV miRNAs, and cfDNA methylation), and models were optimized using hyperparameter tuning. The nested CV framework allowed for unbiased selection of features and performance metrics, mitigating overfitting and ensuring reproducibility. The primary goal was to determine the sensitivity of each modality in identifying advanced adenomas (AA) and colorectal cancer (CRC) at a fixed specificity of 91%, a threshold selected to reflect clinical relevance in screening settings, based on US Centers for Medicare and Medicaid Services (CMS) guidelines. For each omics modality, models were trained to distinguish positive cases (AA + CRC) from negative controls (Healthy + Non-AA), and sensitivity was calculated on held-out test sets during each iteration of the cross-validation. Model performance varied across omics.

EV gene expression emerged as the most sensitive modality. At 91% specificity, sensitivity for detecting advanced adenomas was approximately 54.8% (95% CI 26.4%-75.6%), while sensitivity for CRC was 94.1% (95% CI 79.2%-100%). Correspondingly, the AUROC for AA detection using EV gene expression averaged 0.81, while CRC detection achieved an AUROC of 0.96. These metrics suggest strong discriminatory power and potential for clinical deployment. EV miRNAs and proteomics showed intermediate performance. Sensitivity for AA detection for EV miRNA and proteomics were 41.5% (95% CI 19.6%-60.6%) and 30.0% (95% CI 13.9%-50%), with AUROC values of 0.71 and 0.70, respectively. CRC detection was more reliable, with sensitivities for EV miRNA and proteomics at 76.8% (95% CI 51.7%-92.4%) and 64.2% (95% CI 43.6%-87.2%), with AUROC values of AUROC 0.93 and 0.91, respectively. These modalities may provide complementary information but were less effective than gene expression alone (Figures 5a-d).

**Figure 5.**
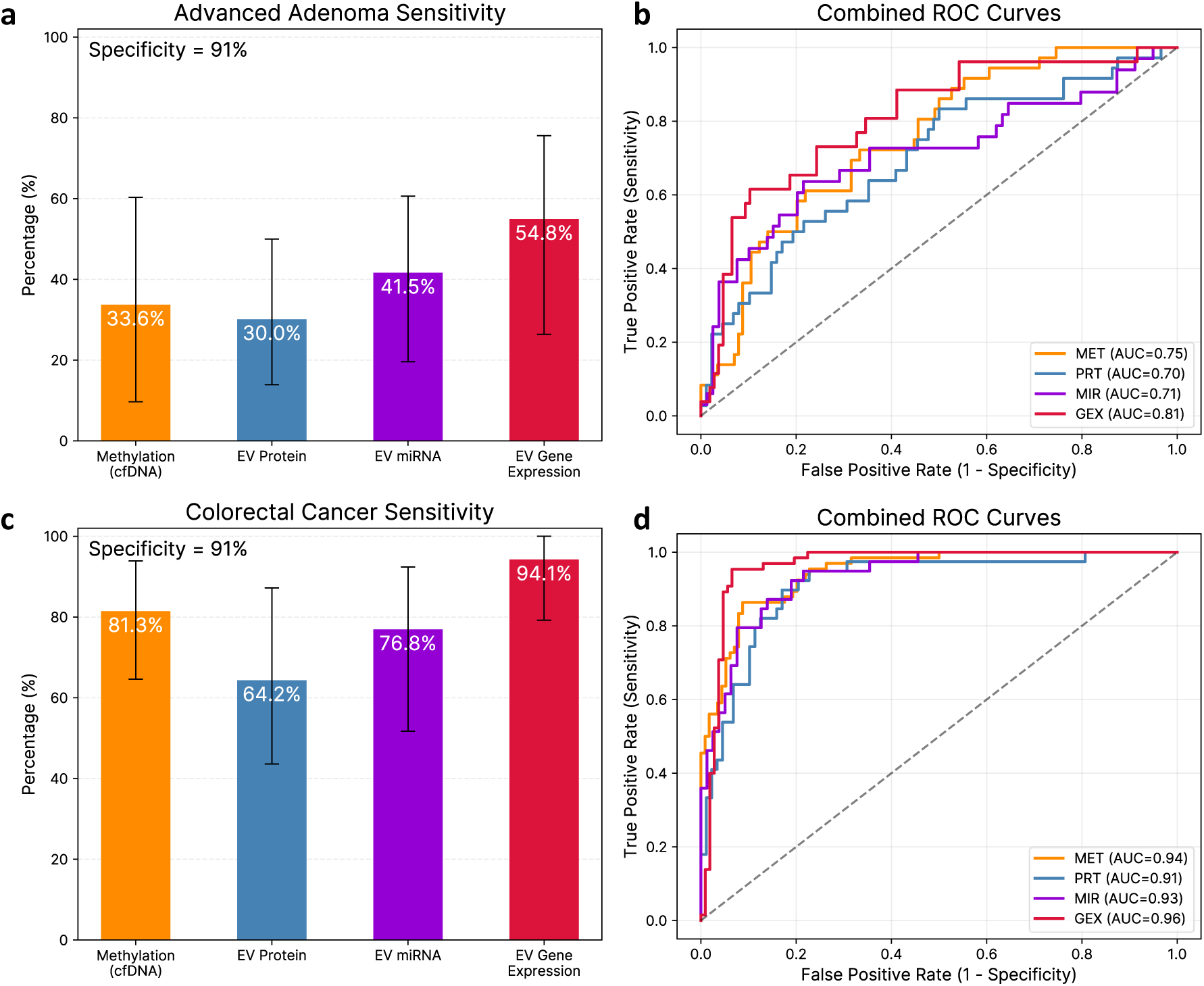
Performance of models under cross-validation indicates high sensitivity to pre-cancerous lesions from EV gene expression. Models were trained using a maximum of 20 features from each omic datatype to distinguish Healthy+Non-AA from AA+CRC and evaluated on held-out samples of each specific condition being assessed. **a.** Sensitivity for detecting advanced adenomas at 91% specificity across different omics platforms. **b.** Aggregate AUROC curve for advanced adenoma detection from all predictions across 5-fold cross-validation with 10 repeats. **c.** Sensitivity for detecting colorectal cancer (stages 1-4) at 91% specificity across different omics platforms. **d.** Aggregate AUROC curve for colorectal cancer detection (stages 1-4) from all predictions across 5-fold cross-validation with 10 repeats.

While widely used in commercial assays^24,25^, cfDNA methylation demonstrated lower sensitivity in this cohort. Detection of AA was lower at 33.6% (95% CI 9.7%-60.3%), and CRC sensitivity was 81.3% (95% CI 64.6%-93.9%), suggesting limited utility for early-stage detection in plasma. These findings align with prior literature indicating that cfDNA is less abundant and less informative in early disease stages.^26^

To visualize performance, ROC curves were plotted for each modality and condition. EV gene expression curves showed steep rises and high true positive rates, confirming superior performance. In contrast, cfDNA methylation curves were flatter, indicating reduced sensitivity at high specificity thresholds (Figures 5b, 5d).

In summary, EV gene expression demonstrated the highest sensitivity and AUROC for detecting both pre-cancerous and cancerous lesions. While other modalities provided reasonable signals, none matched the performance of gene expression alone. These results underscore the value of EVs as a source of circulating biomarkers and support their use in non-invasive screening for colorectal neoplasia.

### Multiomic Signature Reveals a Molecular Continuum in Colorectal Cancer Progression

To identify the most robust and biologically relevant biomarkers from our multiomic analysis, we investigated the features most frequently selected during cross-validation. Figure 6 provides a visual representation of the relative abundance of features selected in more than three validation folds, illustrating their expression patterns across healthy, Non-AA, AA, and CRC patient groups. The data are presented on radar plots where feature levels in the AA and CRC cohorts are normalized to the average of the healthy control cohort, which is represented by the blue unit circle baseline. This method effectively highlights the magnitude and direction of dysregulation for the most diagnostically powerful features, revealing a clear continuum of molecular alterations during the progression of colorectal neoplasia.

**Figure 6.**
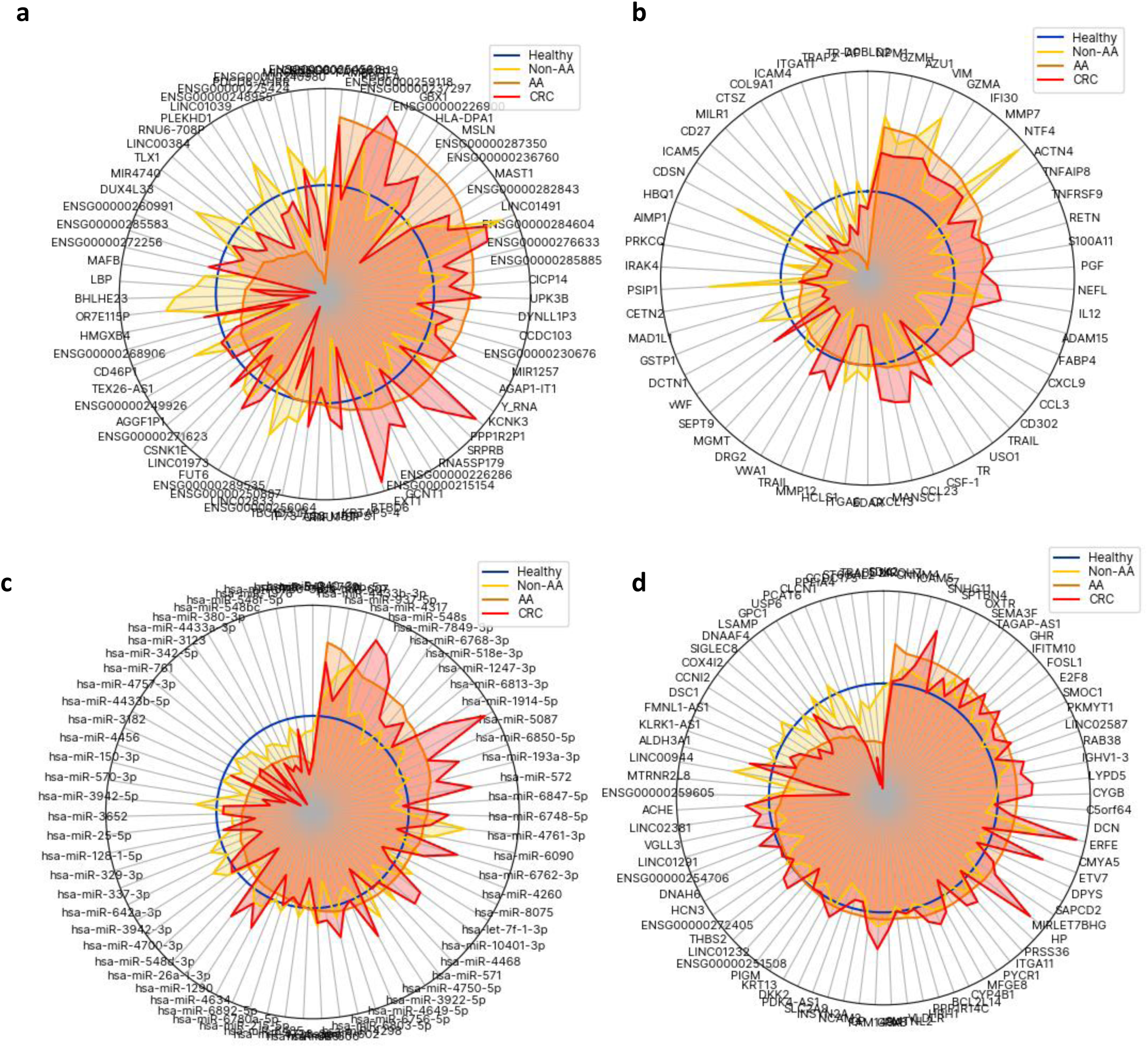
Representation of features selected more than three times during cross-validation. Features are normalized to average Healthy feature quantities (blue circle) as described in Methods. Points inside the blue circle have lower expression/feature quantity than the healthy equivalent, while those outside the blue cirvle have higher expression/feature quantity. **a**. Methylation beta values by condition (76 features selected). Features are labeled by the starting coordinate **b.** Proteomic expression by condition (58 features selected). **c.** miRNA expression by condition (71 features selected). **d**. Gene expression by condition (81 features selected).

The analysis revealed distinct and biologically significant patterns of dysregulation across the four omic layers, corroborating known mechanisms of colorectal carcinogenesis. In the methylation panel, many consistently selected genes demonstrated progressive hypermethylation from healthy controls to AA and finally to CRC, with cancer showing the most profound changes (Figure 6a). This pattern of hypermethylation is a well-established hallmark of CRC.^27^ In addition, the methylation data identified locus-specific hypermethylation of FUT6 (Fucosyltransferase 6), which was elevated in both AA and CRC samples (Figure 6a). FUT6 has previously been identified as a significant protective biomarker in head and neck squamous cell carcinoma (HNSCC), with its expression notably reduced in cancerous tissues compared to normal ones^28^; higher FUT6 levels correlate with improved overall survival in patients. Functional studies demonstrate that FUT6 inhibits key tumor cell behaviors such as proliferation, migration, invasion, and epithelial-mesenchymal transition, underscoring its role in suppressing tumor progression.^29^ In the proteomics data, we observed dysregulation of several key proteins with established roles in CRC (Figure 6b). For instance, CXCL13 (C-X-C motif chemokine ligand 13), was substantially upregulated in both CRC and AA compared to healthy, consistent with existing research demonstrating that elevated CXLC13 expression in the tumor microenvironment promotes CRC progression and metastasis, partly by recruiting tumor infiltrating B lymphocytes, and is associated with poorer patient prognosis.^30^ In parallel protein SEPT9 (Septin 9), involved in cytokinesis, and cell cycle control, was downregulated in AA and more significantly in CRC compared to healthy, consistent with reports linking its known hypermethylation in colorectal, and is an FDA-approved biomarker for CRC screening^31^(Figure 6b). The miRNA and gene expression data further reinforced these findings, identifying critical regulators of cancer pathways (Figure 6c, 6d). This theme of oncogenic dysregulation was echoed in the regulatory and transcriptional data. Among the selected miRNAs, hsa-miR-1914-5p was significantly upregulated in CRC (Figure 6c), consistent with its emerging role as an “onco-miR” that promotes colorectal cancer cell proliferation and metastasis by targeting tumor suppressor genes.^32^ Finally, the gene expression analysis identified a marked increase in the transcript for *ETV7* (Ets Variant Transcription Factor 7), particularly in the CRC state (Figure 6d). As an oncogenic transcription factor, *ETV7* can promote cell survival and tumor progression, and its overexpression has been associated with cell proliferation, migration, and cell cycle amplification.^33^ The gene expression analysis also revealed the overexpression of *ERFE* (Erythroferrone) in both AA and CRC. *ERFE* contributes to the progression and invasion of colorectal cancer (CRC) through its involvement with TP53 mutations and its participation in key signaling pathways like PI3K/AKT and mTOR.^34,35^ It may also activate the NOTCH-related signaling pathway.^36^ Additionally, its overexpression is associated with a poor prognosis and may play a role in the tumor’s ability to evade the immune system.^37^ Taken together, the consistent selection of these specific biomarkers demonstrates that our models have independently identified key drivers of CRC pathogenesis across multiple biological layers.

## Discussion

### Extracellular Vesicle Gene Expression as a Superior Modality for Colorectal Neoplasia Detection

This study conducted a comprehensive, multiomic comparison to identify the most promising circulating biomarkers for the early detection of colorectal neoplasia. Our central finding is the superiority of EV-derived gene expression over other analytes, including cfDNA methylation, EV proteomics, and EV small RNAs. The EV gene expression signature not only achieved the highest sensitivity for both AA and CRC but also revealed biologically coherent pathway alterations that mirror the known progression of the disease.

Our machine learning models, rigorously evaluated using a nested cross-validation framework, demonstrate that EV gene expression achieved a sensitivity of 54.8% for AA and 94.1% for CRC, at a clinically relevant specificity of 91%. This performance surpasses that of other modalities; for instance, cfDNA methylation showed a lower sensitivity of 81.3% for CRC and 33.6% for AA in our cohort. The robustness of the EV-RNA signal can be attributed to the biology of EVs, which act as natural protectors of their molecular cargo, shielding RNA from degradation by circulating RNases.^38^ This is a critical advantage over cell-free RNA, which is notoriously unstable in plasma.^39^ Our tissue-of-origin deconvolution analysis supports this, identifying a substantial and consistent contribution from the colon to the plasma EV-RNA pool.

Crucially, this superior performance is rooted in the underlying biology of CRC. Our unsupervised pathway analyses confirmed the progressive enrichment of key cancer hallmark pathways, such as Apoptosis and Epithelial-Mesenchymal Transition (EMT), across the healthy-adenoma-carcinoma sequence. The ability to detect these transcriptional programs within circulating EVs suggests this liquid biopsy approach provides a real-time, systemic snapshot of active tumor biology. This contrasts with cfDNA, which is often considered a product of apoptosis and necrosis, representing remnants of dead cells.^40^ The active shedding of EVs by living tumor cells, carrying cargo that reflects ongoing transcriptional programs, may offer a more dynamic biomarker for detection and monitoring therapeutic response.

### EV Gene Expression Outperforms Combined Omic Approaches

A natural next step to building a signature was to evaluate if integrating multiple omics modalities would yield superior test performance compared to using single omics alone. Counterintuitively, our preliminary results show that EV gene expression alone outperformed all other modalities and that a combined model offered no significant improvement. This prompts a deeper analysis into the nature of the information provided by each omic layer (data not shown).

To understand this result, we used Multiple Co-Inertia Analysis (MCIA)^41^ to explore the underlying relationships between the datasets. The analysis revealed a striking pattern: while the different omic signals were highly concordant in healthy and Non-AA samples, they showed extreme molecular divergence with the progression to AA (Figure S6a) and CRC (Figure S6b). In the MCIA projection, individual CRC samples exhibited long vectors, indicating that each omic layer was capturing distinct and dramatic shifts specific to the cancerous state. This divergence explains why a simple combination of omics did not improve classification—the signals are not merely redundant but are highly complementary. While correlated in health, each omic layer reveals unique facets of dysregulation during carcinogenesis. This underscores the value of our multi-omic approach; although a single modality may be “fit-for-purpose” for one classification task, capturing this complex and heterogeneous landscape is essential for a truly comprehensive understanding of CRC and holds significant promise for developing more advanced models in future, larger cohort studies.

One apparent paradox was the performance of EV-derived miRNAs. While unsupervised analysis showed a strong biological signal (large average fold change), the supervised classification model yielded lower sensitivity (30.0% for AA, 78.9% for CRC) than the EV gene expression model. Our findings indicate that the superior performance of the mRNA-based model is attributable to the high-dimensional nature of the data input, even when employing feature selection to derive a more parsimonious final model. The vast transcriptome provides a rich feature space from which a limited subset of highly informative mRNAs can be selected, yielding a clear and separable signature with potent classification power. This contrasts with the signal derived from miRNAs, which, despite their established stability as biomarkers, exert their influence by perturbing complex and interwoven regulatory networks.^42^ This likely reflects the pleiotropic nature of miRNAs; a single miRNA can modulate hundreds of target genes^43^, creating a powerful but functionally diffuse biological effect, making it inherently more challenging to isolate a comparably specific set of features. Our data suggests that for this specific classification task, the direct, high-dimensional information content of the EV transcriptome is more valuable for achieving high sensitivity and specificity.

### Methodological Rigor and Critical Evaluation of Study Limitations

The credibility of biomarker discovery studies hinges on methodological robustness and transparent acknowledgment of limitations. A cornerstone of our analytical approach was the use of a nested cross-validation (CV) framework. This is the standard for biomarker discovery, as it provides an unbiased estimate of model performance by strictly separating model tuning from performance evaluation, thereby preventing information leak and overly optimistic results that fail to generalize.^44^ Furthermore, as a multi-site study, our analysis pipeline incorporated computational batch correction methods (ComBat-seq^45^, OLS corrections) to mitigate non-biological variation arising from differences in sample handling, ensuring that the observed signals are biological, not technical artifacts.

Despite these strengths, our study has important limitations rooted in our cohort’s composition. One of them is the insufficient representation of lesions from the serrated pathway of carcinogenesis. This pathway, often initiated by BRAF mutations and characterized by a CpG Island Methylator Phenotype (CIMP), accounts for up to 30% of all CRCs and includes precursors like sessile serrated lesions (SSLs).^46^ Similarly, though all CRC stages were represented in the study cohort, it was heavily weighted towards the more advanced disease states (stages III-IV) compared to large screening cohort population studies (see Table 1).^47^ An effective screening test must detect precursors from both the traditional and serrated pathways. Future validation studies must ensure the dedicated recruitment of patients with well-characterized serrated lesions to evaluate performance against this distinct biology.

Furthermore, the lack of ethnic diversity in our cohort (92.2% Caucasian) is a limitation. Significant evidence documents racial and ethnic disparities in CRC biology, incidence, and mortality.^48^ Black Americans, for example, have higher incidence and mortality rates and different underlying molecular profiles.^49^ A biomarker developed almost exclusively in a Caucasian population may not perform equivalently in other groups, potentially exacerbating existing health disparities. Therefore, ensuring robust diversity in all future validation cohorts is not just a scientific best practice; it is an ethical imperative for developing equitable healthcare technologies.

### Path to Clinical Translation and Future Directions

This study provides a strong rationale for advancing EV-derived gene expression as a non-invasive test for colorectal neoplasia. The independent identification of biologically relevant features—such as the downregulation of SEPT9 protein, the upregulation of the oncogenic transcription factor *ETV7*, and the increased expression of the onco-miR hsa-miR-1914-5p—underscores the validity of our approach. Our next steps towards clinical implementation require significant investment of resources including running of large-scale, multi-center, prospective validation study in a diverse and representative screening population, including patients with all relevant precursor lesions.

In conclusion, this study demonstrates the superior potential of plasma EV-derived gene expression as a non-invasive biomarker for colorectal neoplasia. By capturing a rich, biologically relevant transcriptomic signal, this modality appears to overcome the limitations of other analytes. While the path forward is challenging, EV-based liquid biopsies hold the transformative promise of improving CRC screening, enabling earlier detection, and ultimately reducing the global burden of this disease.

---

**Figure S1.**
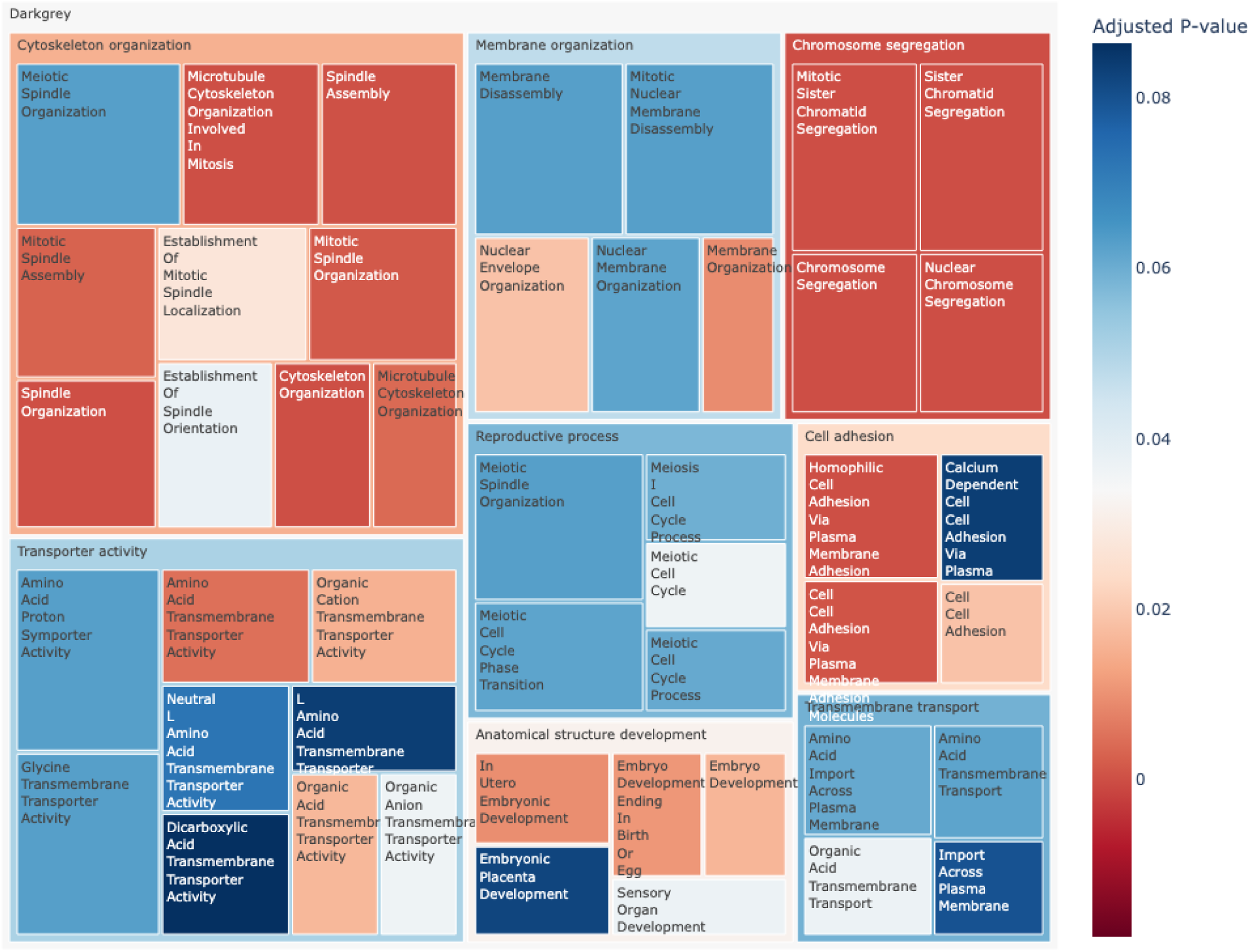
CRC Module. Treemap of gene ontology (GO) pathway enrichment results for the Darkgrey module (positively correlated with Stage 1 or later CRC). GO pathways are categorized at a higher level to summarize information. Box sizes represent combined score combining p-value and odds ratio.

**Figure S2.**
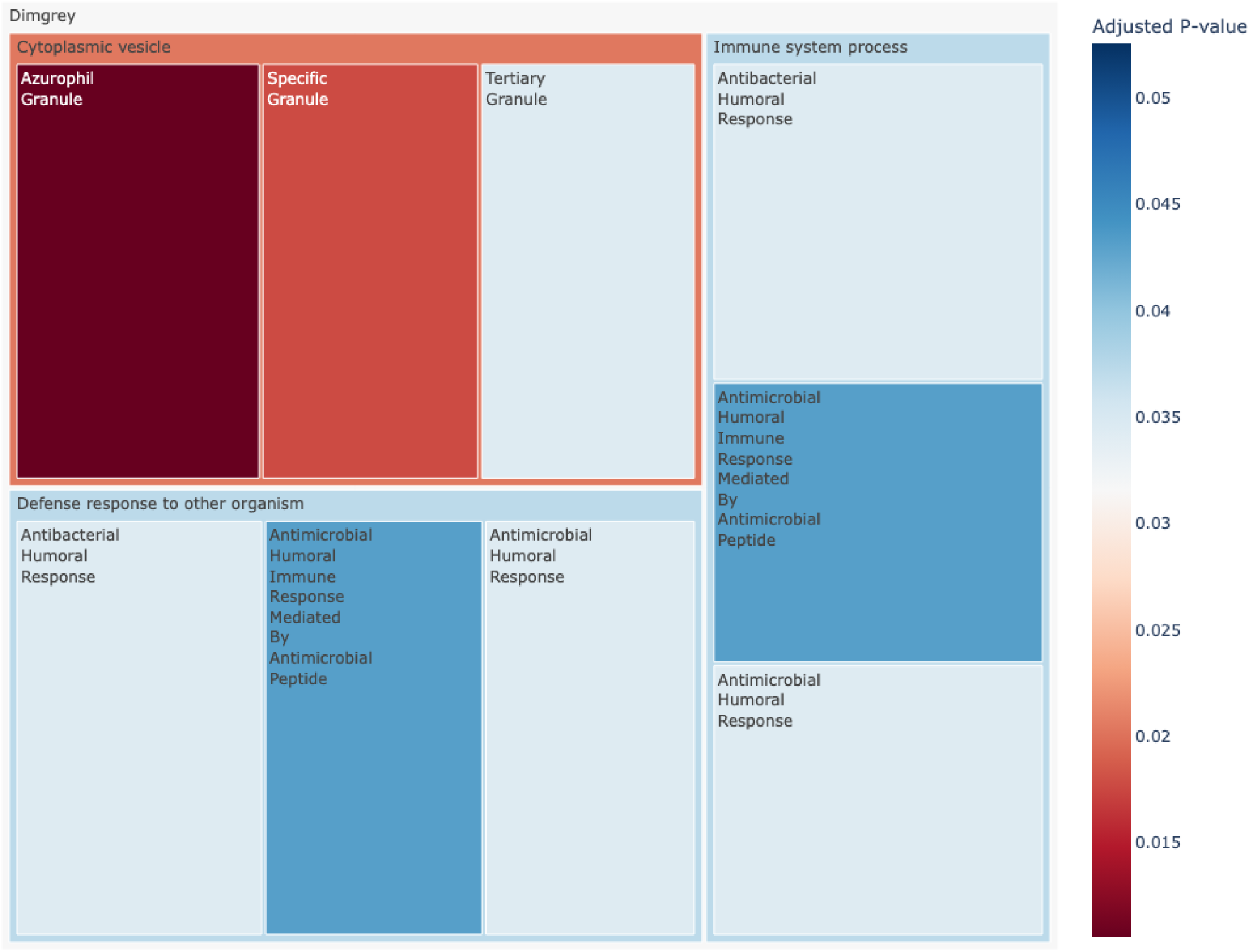
AA module. Treemap of gene ontology (GO) pathway enrichment results for the Dimgrey module (positively correlated with the advanced adenoma condition). GO pathways are categorized at a higher level to summarize information. Box sizes represent combined score combining p-value and odds ratio.

**Figure S3.**
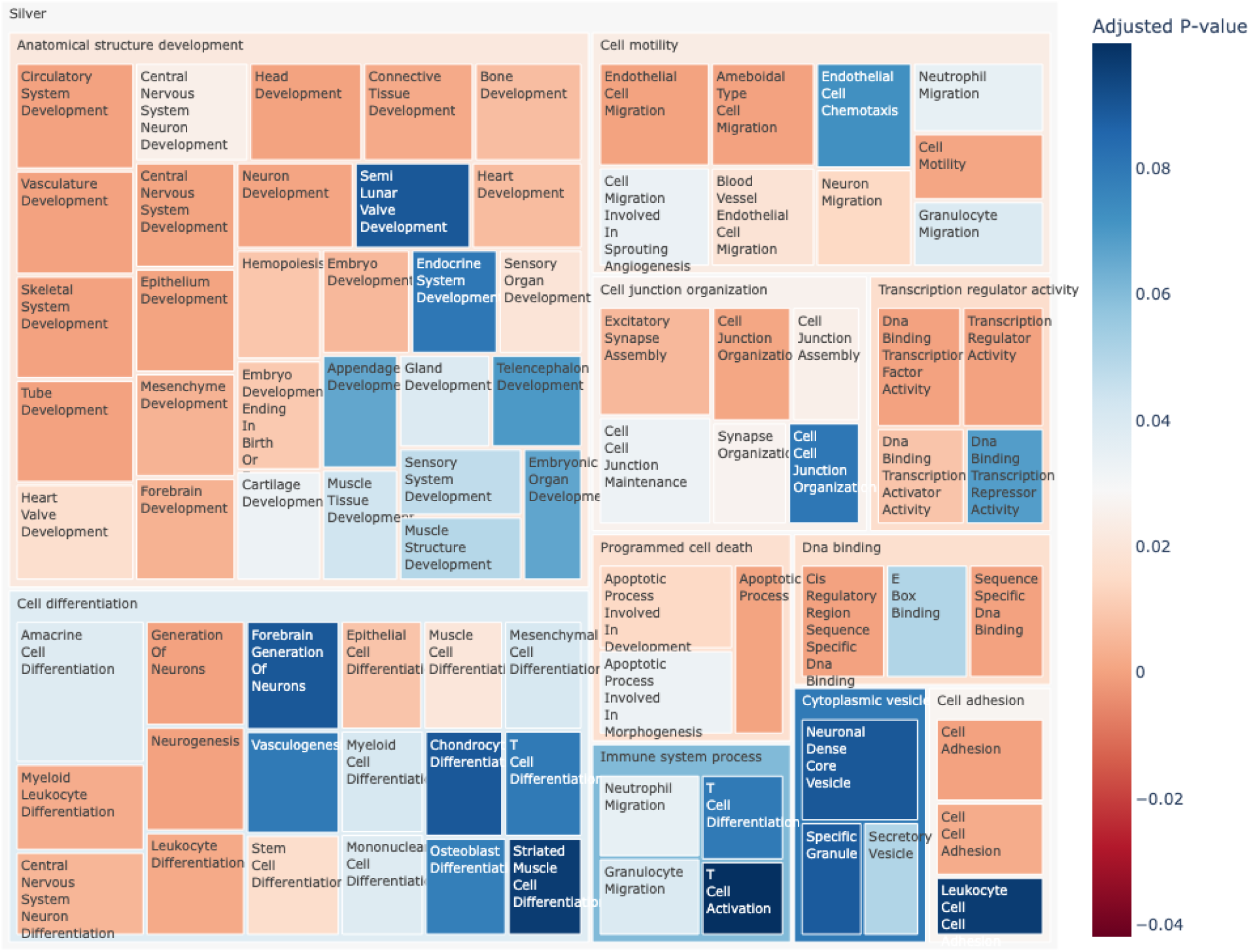
AA/CRC module. Treemap of gene ontology (GO) pathway enrichment results for the Silver module (positively correlated with advanced adenoma and negatively correlated with Stage 1 or later CRC). GO pathways are categorized at a higher level to summarize information. Box sizes represent combined score combining p-value and odds ratio.

**Figure S4.**
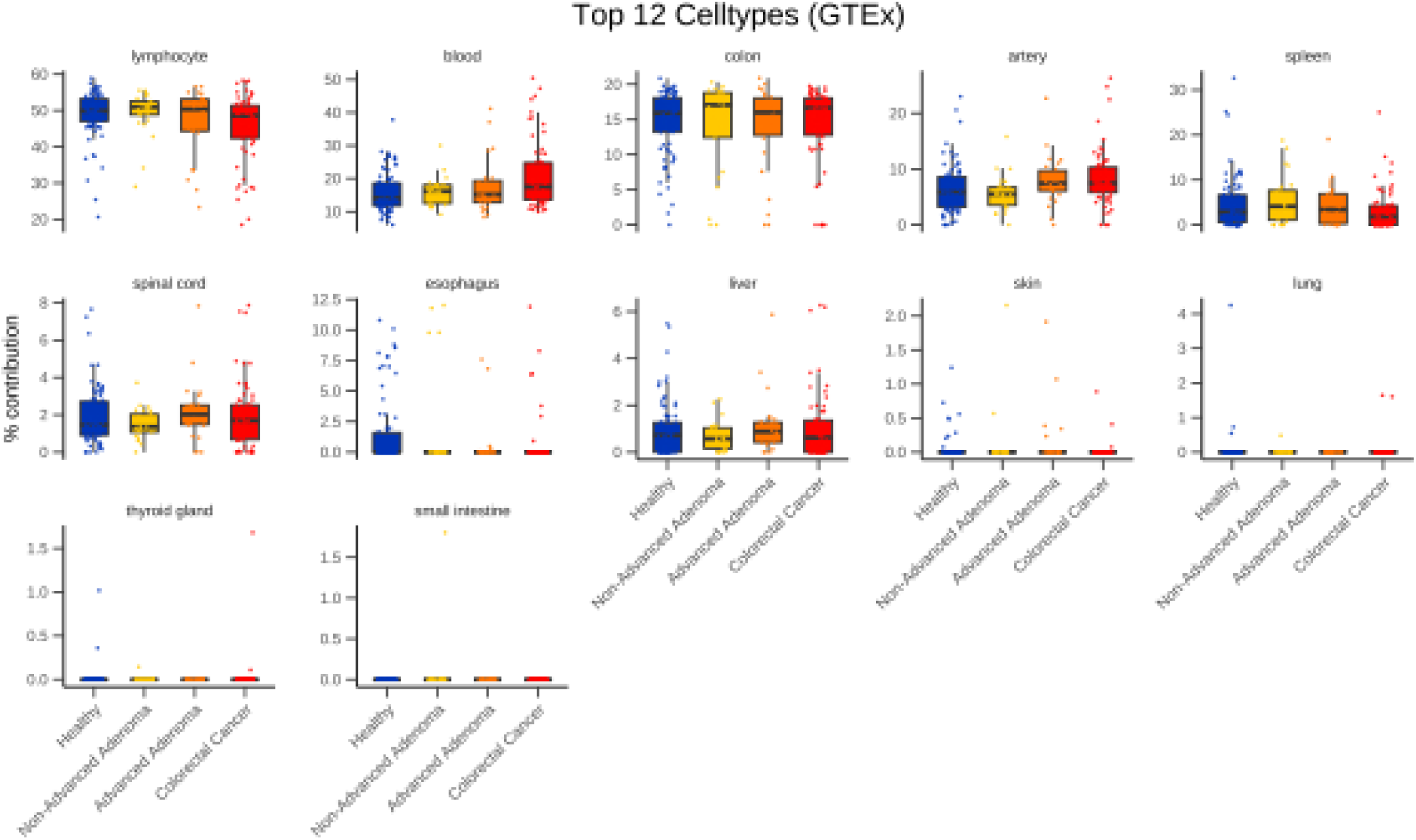
Cell Type Deconvolution. Cell type composition analysis across sample groups for the top 12 most abundant cell types. Cell type composition was estimated from plasma EV-RNA-seq data using the DeconRNASeq algorithm and reference gene expression data from GTEx Consortium. Of the 32 tissues represented in the reference signature matrix, PCA results indicated that 12 cell types were identified in the sample mixtures. Estimated composition contributions are shown for the 12 tissue cell types, grouped by condition. Box plots display the median (center line), interquartile range (box), and whiskers extending to 1.5x the interquartile range. Individual data points are overlaid as points. The y-axis represents the estimated cell type proportion (percent). Contributions from lymphocytes and blood are the largest, as is expected from the most abundant circulating cells contributing EVs to plasma. Notably, contribution from colon (∼15%) is third highest, reflecting the significant role of gastrointestinal tract in systemic EV circulation. Statistical differences between conditions were assessed using one-way ANOVA for each cell type, with p-values adjusted for multiple testing using false discovery rate (FDR) correction. The only cell type with significant differences after correction was ‘blood’ (P_adj_ = 0.021).

**Figure S5.**
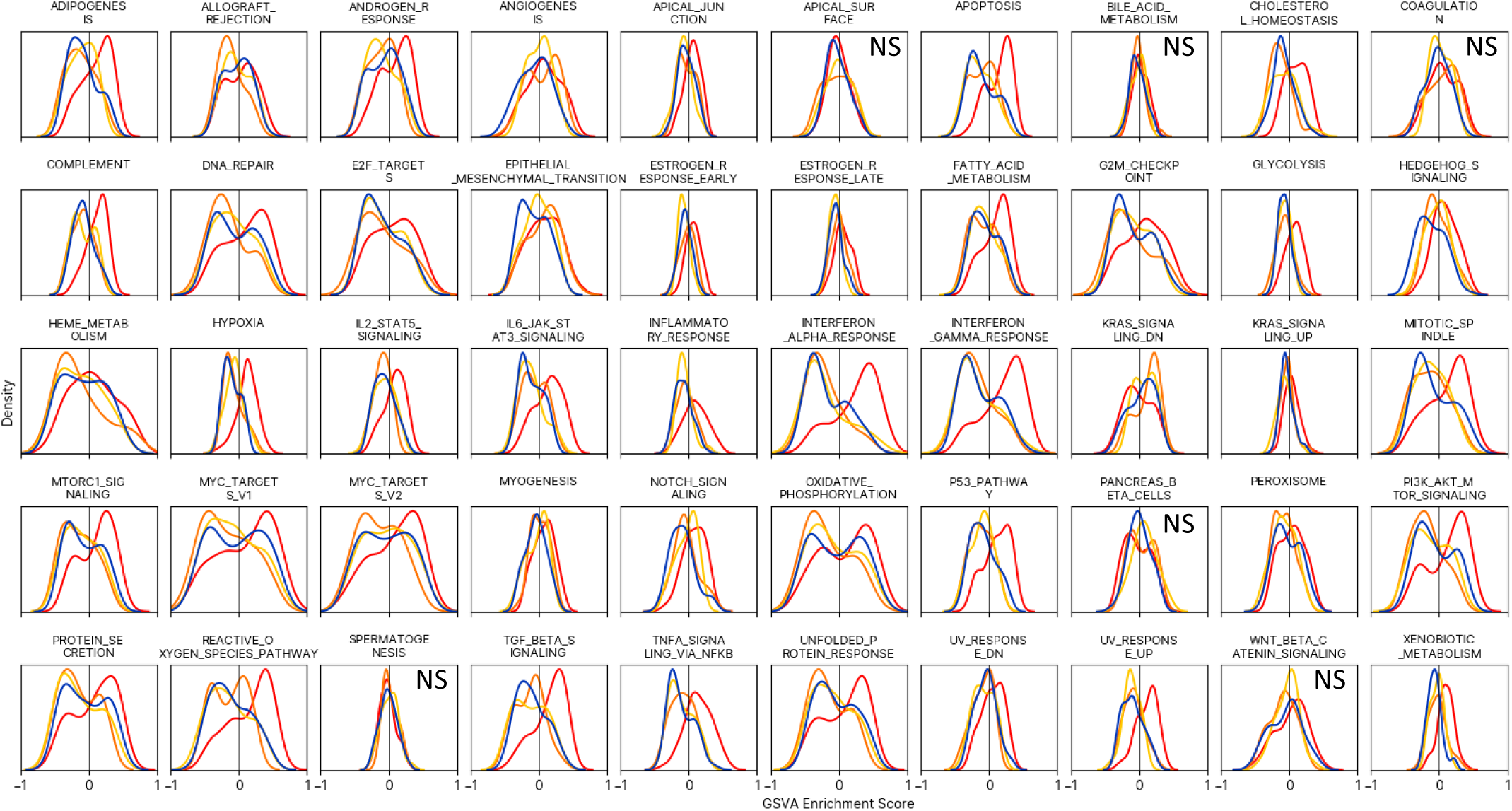
Pathway enrichment. Hallmark pathway enrichment score distributions from gene set variation analysis. All are significant (ANOVA FDR<0.05) unless noted.

**Figure S6.**
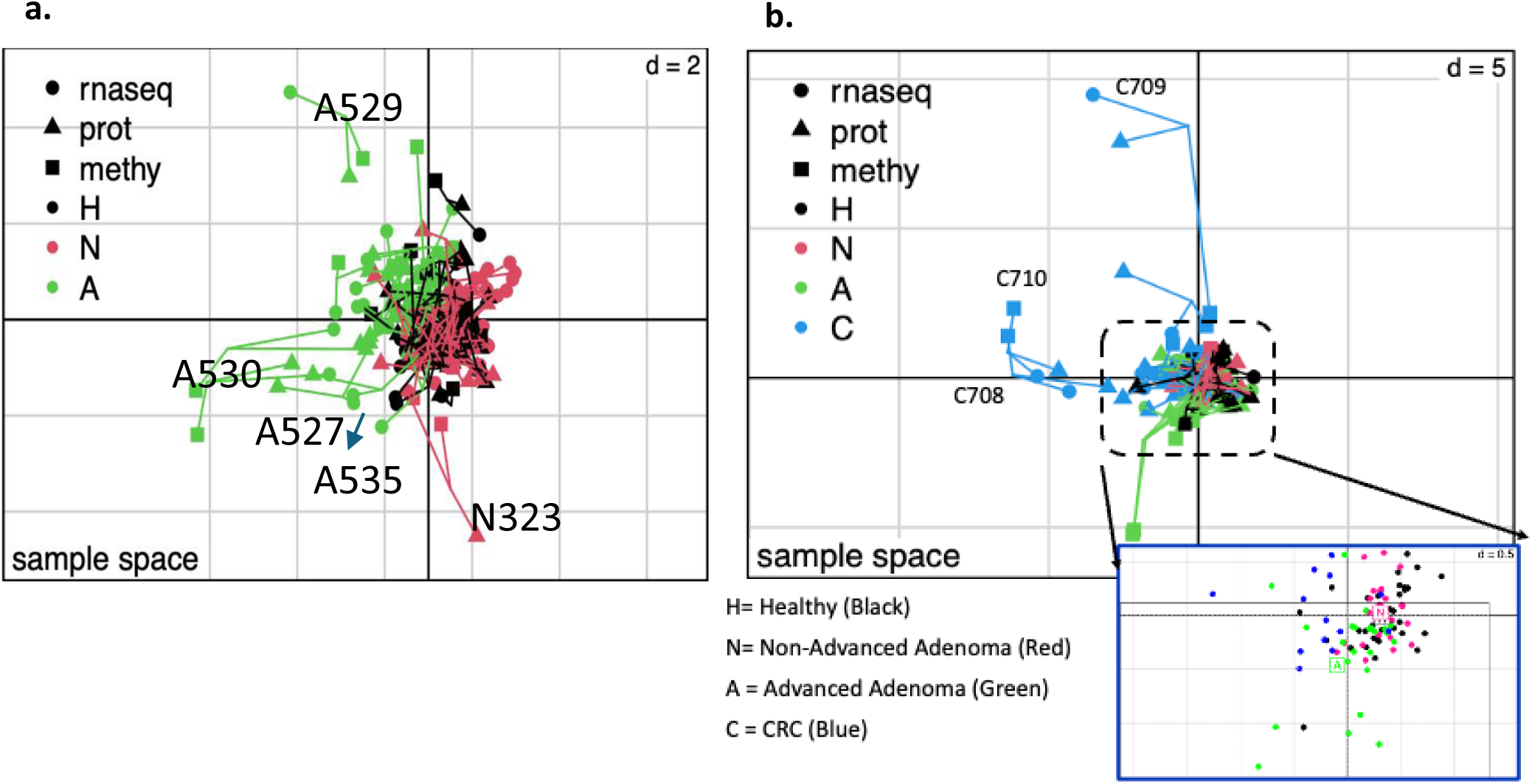
Multi-omics coinertia Analysis. a. Multiple Co-Inertia Analysis of multi-omic data separates Advanced Adenoma (A, green) samples from the tightly clustered Healthy (H, black) and Non-Advanced Adenoma (N, red) groups. The increased spread within the A cluster indicates significant molecular dysregulation. b. The inclusion of Colorectal Cancer (CRC, blue) samples demonstrates extreme molecular divergence. The long lines connecting the omics for CRC samples signify profound molecular alterations, causing them to separate distinctly from all other groups.

