## Supplementary material for "Extracellular Vesicle Gene Expression Enables Sensitive Detection of Colorectal Neoplasia": Methods

Materials and Methods

**Study Design and Population**

This study aimed to identify single and multiomic signatures distinguishing nominally healthy (H) individuals and patients with non-advanced adenomatous polyps (NA) from those with advanced adenomas (AA) and colorectal cancer (CRC) stages I-IV using extracellular vesicle (EV) multiomics (transcriptomics and proteomics) and cell-free DNA (cfDNA) methylomics derived from plasma liquid biopsy samples. Plasma samples were collected prior to colonoscopy from participants who underwent screening as part of this study. Participants were categorized into four disease classification cohorts (H, NA, AA, CRC) based on colonoscopy results and histological evaluation. Non-advanced adenoma is defined as any adenomatous lesion that was not advanced adenoma or colorectal cancer and that does not have serrated lesions. Advanced adenoma is defined as an adenomatous lesion that: (i) has histologically proven dysplasia or/and (ii) is ≥10 mm large or/and (iii) has a villous or tubulovillous component or is serrated. Advanced adenoma is defined as having (i) histologically proven (low or medium/high grade) dysplasia and/or (ii) has villous or tubulovillous component and/or has serrated lesions. Cancer Stages were classified according to AJCC (I-IV).

**Sample Collection and Processing**

This was a prospective multi-center study conducted across 5 sites (two US, 3 ex-US). Eligibility criteria required that participants be classified by a physician or healthcare provider as being at average risk for CRC. Individuals with a positive fecal immunochemical test (FIT) or fecal occult blood test (FOBT) result within the previous 6 months were excluded. Recruitment was conducted among individuals scheduled to undergo routine CRC screening, including colonoscopy, at participating clinical centers. Written informed consent was obtained from all participants prior to enrollment. The study protocol was approved by the Institutional Review Boards of all participating institutions. Whole Blood samples were collected in K2EDTA tubes only. Plasma samples were prepared from 20 – 40 (2-4 x 10mL) of blood. Plasma was processed within two hours of collection. A minimum volume of 10 mL plasma was collected per subject. Samples were aliquoted into 1.5-2ml cryovials and stored at -80°C until further processing.

**Wet Lab Methods**

**EV isolation & RNA extraction:**

EV isolation and Total RNA extraction was performed on 2 mL of filtered plasma using Exosome Diagnostic’s proprietary ExoLution workflow. RNA was assessed for quality using Agilent Bioanalyzer with the RNA Pico 6000 Kit (Agilent Technologies). Isolated RNA was stored at -80 °C prior to library generation.

**Long RNA Profiling**

Profiling of long RNA was performed using Exosome Diagnostics’ proprietary EV long RNA sequencing method. Briefly, isolated RNA from ExoLution method described above was treated with DNase to remove any co-purified DNA present in the sample. Synthetic RNA spike-in controls (ERCC spike-in mix, Thermo Fisher Scientific) were then added to each sample. Total RNA was then fragmented and reverse transcribed. Adapter addition was performed using a PCR-based approach. Libraries were purified and size selected via two rounds of AMPureXP® beads (Beckman Coulter). The final libraries were then amplified by PCR, followed by clean up using AMPureXP® beads. Libraries were quantified using the Qubit 1X dsDNA HS Assay Kit (Thermo Fisher Scientific). Equimolar amounts of each library were pooled in batches of eight for hybrid capture using manufacturer’s protocol (Agilent Technologies). Libraries were hybridized with custom bait sets covering ERCC spike-ins, all protein coding genes and long noncoding RNAs (hg38). Hybridized libraries were captured, washed and amplified. Pooled libraries were quantified using the Qubit 1X dsDNA HS Assay Kit (Thermo Fisher Scientific) and assessed for size distribution using the TapeStation High Sensitivity D1000 or D5000 tape kit (Agilent Technologies). Final libraries were pooled and sequenced on Illumina® NextSeq2000 using 2 × 151 cycles read length chemistry with PhiX control.

**Small RNA Profiling**

Profiling of small RNA was done using Exosome Diagnostics EV small RNA sequencing method, based on RealSeq Biosciences biofluids small RNA kit, cat. 600-00048. Briefly, isolated RNA was treated with DNase to remove any co-purified DNA present in the sample. A synthetic miRNA spike-in control (QiaSeq miRNA spike-in, cat. No. 331535, Qiagen) was added to the samples. 3-prime adapters were added to the small RNA using a ligation-based approach. The ligated RNA was then circularized. Following removal of dimers using SPRI beads, the samples were reverse transcribed. Final libraries were amplified using a PCR-based approach. Libraries were then cleaned up using AmpureXP beads, and size selected for enrichment of all transcripts from 15-200 nucleotides in length using the PippinHT (Sage Science). Size-selected libraries were quantified using the Bioanalyzer High Sensitivity DNA (Agilent Technologies) and the Qubit 1X dsDNA HS Assay Kit (Thermo Fisher Scientific). Final libraries were pooled and sequenced on Illumina® NextSeq2000 using 1x151 cycles (single end) read length chemistry with PhiX control.

**cfDNA Isolation**

cfDNA extraction was performed on 2 mL of filtered plasma using Exosome Diagnostics’ proprietary ExoLution Plus workflow. cfDNA was assessed for residual genomic DNA using the TapeStation cfDNA kit (Agilent Technologies). Samples with a ratio of greater than 2:1 gDNA: cfDNA underwent high molecular weight clean up using AMPureXP® beads (Beckman Coulter). Cleaned up samples were reassessed for gDNA contamination using the TapeStation. Isolated cfDNA was stored at -80 °C prior to library generation.

**Methylation Profiling**

cfDNA methylation profiling was performed using a modified protocol based on New England Biolab’s NEBNext Enzymatic methyl-seq kit (cat. No. 101977, Twist Biosciences). Briefly, a zero percent methylated reference DNA and a one hundred percent methylated reference DNA were fragmented using the Covaris M220 Focused Ultrasonicator and added to isolated cfDNA as internal sample controls. cfDNA underwent end repair, followed by ligation of sequencing adapters. Samples then went through enzymatic conversion of unmethylated cytosines to uracils. Unique dual identifiers were added using PCR. Libraries were quantified using the Qubit 1X dsDNA HS Assay Kit (Thermo Fisher Scientific). Equimolar amounts of each library were pooled in batches of eight for hybrid capture using manufacturer’s protocol (Twist Biosciences). Briefly, libraries were hybridized with bait sets against a Human Methylome Panel (cat. No. 105520, Twist Biosciences). Hybridized libraries were captured, washed and amplified. Pooled libraries were quantified using the Qubit 1X dsDNA HS Assay Kit (Thermo Fisher Scientific) and assessed for size distribution using the TapeStation High Sensitivity D1000 or D5000 tape kit (Agilent Technologies). Final libraries were pooled and sequenced on Illumina® NextSeq2000 using 2 × 151 cycles read length chemistry with PhiX control.

**EV Proteomics**

One milliliter of filtered plasma samples was used for extracellular vesicle (EV) isolation using a modified protocol of ExosomeDx’s proprietary Exolution workflow. Purified EVs underwent buffer exchange to phosphate-buffered saline (PBS) using Zeba 7K MWCO columns as per the manufacturer's instructions. For EV lysis, buffer-exchanged EVs were mixed with lysis buffer at a 1:1 ratio and incubated for 15 minutes at room temperature with rotation. Following this protocol, the samples were prepared for downstream Olink analysis or stored at -80 °C for future use. EV proteome was profiled using six Olink Target 96 panels including Immuno-Oncology, Immune Response, Cardiovascular III, Neuro Exploratory, Oncology II, and Oncology III. One microliter of the lysed EV samples was added into the incubation plate, and the Olink's Target 96 protocol was followed for the subsequent steps of incubation, extension, and detection. The data obtained was initially analyzed using the Olink NPX Signature software to perform quality controls, utilizing the internal controls and external controls provided in the kit. Subsequently, the raw data was exported for additional bioinformatics analysis.

**Bioinformatics Analysis**

**Small and Long RNASeq Analysis**

Transcriptomic analysis of long RNA was performed using a [customized Nextflow long RNA pipeline; nf-core/rnaseq]. Briefly, reads passing QC were trimmed, UMI-deduplicated, and aligned to the human reference transcriptome (Ensembl GRCh38.108) by STAR (v2.7.10a) and quantified by Salmon (v1.9.0).^1,2^ Quality control measures included minimum sequencing depth threshold (≥15 million reads/sample), and mapping quality assessment. Normalized transcripts per million (TPM) counts were used to evaluate technical reproducibility between all samples. To enable in-fold batch correction, CPM of length-normalized readcounts were used as the input to subsequent analysis. A full description of the long RNA-seq pipeline and quality control protocol is in the Supplementary Methods.

Analysis of small RNA was performed using a [customized Nextflow small RNA pipeline; nf-core/smrnaseq]. Briefly, raw reads passing QC filters were adapter trimmed and filtered by minimum size (15 nt). Reads were sequentially aligned (Bowtie, v1.3.0)^3^ against the miRbase reference mature miRNA database (release 22.1) and miRbase reference hairpin sequences (release 22.1).^4^ Post-alignment, TMM (trimmed mean of m values) normalized counts for reads aligned to mature and precursor miRNAs were calculated using edgeR (v4.0.16).^5^ A full description of the small RNA-seq pipeline and quality control protocol is in the Supplementary Methods. Normalized CPM counts of mature miRNAs were used as inputs for feature selection, cross-validation and model performance evaluation in the cross-validation pipeline.

Differential gene expression (DEx) was calculated using a customized Nextflow pipeline for DESeq2 analysis.^6^ Input read counts for DEx were batch-corrected using ComBat-seq^7^  to account for collection site and sequencing flow cell batch effects. Genes identified by differential expression were further analyzed for biological pathway enrichment (Gene Set Variation Analysis, GSVA). High variance genes were passed to WGCNA^8,9^ to identify co-expressed gene modules, which were then further correlated with disease status. It works by first calculating a correlation matrix for all genes and then using it to build a network where genes are nodes connected by edges representing their co-expression strength. The network is then mined to find clusters of densely interconnected genes. These modules, represent sets of genes that are likely functionally related or co-regulated. The final step is to correlate these modules with disease states. Comparisons between plasma EV-derived transcriptomics and colorectal tissue gene expression were performed to assess coherence and biological relevance.

**Proteomics Analysis**

Proteomic profiling utilized 6 different Olink's Target 96 panels, namely, Oncology II, Oncology III, Immuno-Oncology, Cardiovascular III, Immune Response, Neuro Exploratory, encompassing total of 552 unique protein assays. Normalized protein expression (NPX) values were obtained and used for analysis. Limit of detection (LOD) was established based on the correlation between technical replicates included, with a threshold set at a correlation value below 0.9 to ensure reproducibility. Bridging samples were included in each batch to enable data normalization across multiple batches using OlinkAnalyze R package (v3.9.0) (https://olink-proteomics.r-universe.dev/OlinkAnalyze/). Samples receiving QC warnings using Olink NPX Signature software were further evaluated through principal component analysis (PCA) to identify extreme outliers.

**cfDNA Methylation Analysis**

DNA methylation profiling was conducted using enzymatic methyl-seq (EM-Seq). Methylomic analysis of EM-seq data was performed using a [customized Nextflow pipeline for methylation sequencing data; nf-core/methylseq]. Briefly, reads passing quality filters were trimmed and aligned to the human reference genome (hg38) by ‘bwa-meth’ (version 0.2.2)^10^ . Aligned reads were de-duplicated using Picard ‘MarkDuplicates’ (version 3.0.0) and methylation calls extracted with MethylDackel (version 0.6.0) (https://github.com/dpryan79/MethylDackel). All alignment quality control metrics were performed with samtools^11^ , qualimap^12^ , and preseq (https://github.com/smithlabcode/preseq). Quality control included read depth filtering and assessment of methylation conversion efficiency. Enrichment analysis of methylation changes was conducted to evaluate their role in CRC progression and early detection potential.

Reads were extracted at coordinates corresponding to all transcripts in GRCh38 and fraction methylation was averaged to create a feature-by-sample matrix (we refer to these loosely as beta values). This matrix was used to assess sensitivity of gene methylation in cfDNA to detection of AA and CRC conditions.

Differential methylation was estimated by dividing the log2 beta values between conditions.

**Feature selection and cross-validation**

Feature selection and model evaluation for each individual ‘omic modality were conducted using a nested cross-validation framework to prevent data leakage and ensure robust biomarker identification and generalizability. Briefly, the strategy consisted of an outer cross-validation fold for model evaluation and inner fold for hyperparameter tuning and feature selection. In the outer fold, we used repeated stratified 5-fold cross-validation with ten repeats. Preprocessing consisted of sparsity filtering, min-max and size factor normalization. In the inner fold, features were filtered using variance filtering, a false positive rate (FPR) control, a Lasso based feature selection and Boruta feature selection to identify informative predictors. A grid search was performed to select an optimal feature set and tune support vector machine model hyperparameters. The best performing model from the inner loop was re-trained on the full training set and evaluated on the test set of outer loop. Performance was measured using area under the receiver operating characteristic curve (AUC-ROC), sensitivity, and specificity. A thresholding procedure was used to assess the models at predefined specificity of 0.91.

**Biological Significance Evaluation**

To assess the biological relevance of the identified features, pathway analysis and gene set enrichment analysis were conducted. Gene Set Variation Analysis (GSVA) was used to evaluate key colorectal cancer-related pathway scores across patient disease states.^8^  Coherence of differential gene expression between plasma EVs and colorectal tissue was analyzed to validate findings. Protein-protein interaction networks and integrative multi-omic approaches were employed to contextualize findings within known CRC biology. Additionally, a hallmark pathway enrichment analysis was conducted to identify key molecular processes associated with CRC progression.

**Multiple Co-Inertia Analysis (MCIA)**

To integrate the EV-derived transcriptomics, EV protein, and cfDNA methylation datasets, we utilized Multiple Co-Inertia Analysis (MCIA)^13^ , a multivariate method designed to analyze multiple data tables simultaneously. MCIA is an extension of Co-Inertia Analysis (CIA) that identifies axes of maximal shared variance, or co-structure, across several high-dimensional datasets measured on the same samples. This technique allows for the visualization of the relationships between different omic layers and how they collectively contribute to the separation between sample groups. The analysis was performed using the Omicade4 package in the R statistical environment to project the samples into a common space that represents the consensus of the molecular signals.

**Statistical Analysis**

Differential expression of transcriptomics (long RNA-seq counts) was conducted using negative binomial generalized linear models implemented in DESeq2.^6^ Benjamini-Hochberg correction was applied to adjust for multiple testing and control the false discovery rate.

**Covariates Analysis**

Potential covariates were identified *a priori* based on clinical relevance and their potential to confound associations between biomarkers and disease states. Covariates were evaluated for completeness and continuous variables were tested for normality. To identify covariates significantly associated with disease status, univariate analyses were performed for each variable and multivariable covariate selection was performed using backward stepwise selection with entry and stay criteria set at p = 0.20 and p = 0.15, respectively. This approach was implemented using logistic regression models with disease category as the dependent variable.

Important covariates requiring correction (i.e., putatively arising from technical variation) were corrected both in cross-validation (in fold) and globally for analysis of biological relevance as described below.

**Batch Correction**

To control for technical artifacts arising from differences in sample collection site and sequencing batches, an ordinary least square regression approach was designed to remove technical noise from data, while avoiding data leakage between train and test data in nested cross-validation. Such correction was included for all omics that are confounded by site and batches. For biological relevance analysis, specifically for RNAseq (both long and small), a global batch correction with ComBat-seq^8^ was used.

**Principal Component Analysis**

Batch-corrected (if applicable) feature-by-sample matrices were normalized and scaled as appropriate for the omic (gene expression: log2 CPM; methylation: beta value ranging 0-1; proteomics: NPX). Scikit-learn^14^   was used to calculate the first two principal components and their contribution to variance.

**Radar Plots**

Radar plots were created using Matplotlib^15^ . Features selected more than three times during cross-validation were plotted as follows: the log expression data (linear in the case of methylation average beta values) were standard scaled, and the average Healthy condition was subtracted from each condition. The resulting differences schematically represent the signal detected in cross-validation.

**Ethical Considerations**

The study was approved by the institutional review board (IRB) and conducted in accordance with the Declaration of Helsinki. Written informed consent was obtained from all participants before enrollment.

**Data Availability**

Processed data and analytical workflows will be made available upon reasonable request to the corresponding author, in compliance with institutional and ethical guidelines.
